# Measuring Illumina Size Bias Using REcount: A Novel Method for Highly Accurate Quantification of Engineered Genetic Constructs

**DOI:** 10.1101/388108

**Authors:** Daryl M. Gohl, Aaron Becker, Darrell Johnson, Shea Anderson, Bradley Billstein, Shana L. McDevitt, Kenneth B. Beckman

**Affiliations:** University of Minnesota Genomics Center, Minneapolis, MN, 55455, USA; Department of Genetics, Cell Biology, and Development, University of Minnesota, Minneapolis, MN 55455, USA; Vincent J. Coates Genomics Sequencing Laboratory, University of California, Berkeley, CA, 94720, USA

**Keywords:** Next-generation sequencing, DNA library preparation, PCR-free, Illumina, size bias, clustering

## Abstract

Quantification of DNA sequence tags associated with engineered genetic constructs underlies many genomics measurements. Typically, such measurements are done using PCR to enrich sequence tags and add adapters, followed by next-generation sequencing (NGS). However, PCR amplification can introduce significant quantitative error into these measurements. Here we describe REcount, a novel PCR-free direct counting method for NGS-based quantification of engineered genetic constructs. By comparing measurements of defined plasmid pools to droplet digital PCR data, we demonstrate that this method is highly accurate and reproducible. We further demonstrate that the REcount approach is amenable to multiplexing through the use of orthogonal restriction enzymes. Finally, we use REcount to provide new insights into clustering biases due to molecule length across different Illumina sequencing platforms.

## Introduction

Engineered genetic constructs underlie many experimental techniques in genetics and genomics. For example, targeted perturbation of gene function using RNA interference or CRISPR/Cas9 allows for pooled genome-wide genetic screens that can be read-out through next-generation sequencing (NGS) of the small hairpin RNA (shRNA) (Sims et al. 2011; Rodriguez-Barrueco et al. 2013) or synthetic guide RNA (sgRNA) (Wang et al. 2014; Koike-Yusa et al. 2014; Shalem et al. 2014, 2015) constructs, or associated sequence tags/barcodes (Smith et al. 2009). Transposable elements are also commonly used to mutate or otherwise manipulate genetic loci, and similarly enable genome-scale saturation mutagenesis screens in which the transposon-genome junction is measured using NGS (van Opijnen and Camilli 2013). Lineage-tracing (Bhang et al. 2015; McKenna et al. 2016) and connectomics (Kebschull et al. 2016; Peikon et al. 2017) approaches also rely on NGS-based quantification of molecular tags. In all of these approaches, Polymerase Chain Reaction (PCR) amplification is used to enrich for the sequence tags and to add adapters and other functionalities (e.g. sample-specific barcodes) required for sequencing. However, PCR introduces bias into these measurements. Sequence tags comprised of shRNAs, sgRNAs, transposon:genome junctions, or synthetic barcodes, can all differ in primary sequence and biophysical properties, which, along with other variables such as template concentration and PCR conditions, can influence amplification efficiency in unpredictable ways (Aird et al. 2011; Gohl et al. 2016; Strezoska et al. 2012). Adding unique molecular identifiers (UMIs) can mitigate some of this bias, but increases the complexity of both library preparation and analysis (Kivioja et al. 2011; Kinde et al. 2011). Other approaches such as droplet digital PCR (ddPCR) and NanoString analysis can be used to overcome the quantitative inaccuracies associated with measuring engineered genetic constructs, but lack the throughput and resolution afforded by NGS (Geiss et al. 2008; Hindson et al. 2011).

We have developed a novel method, REcount (**R**estriction **E**nzyme enabled **count**ing) for quantifying sequence tags associated with engineered genetic constructs that is straightforward to implement and allows for direct NGS-based counting of a potentially enormous number of sequence tags. In this approach, an Illumina adapter-flanked DNA barcode is liberated by digesting with *M/y*I (a type IIS restriction enzyme that produces blunt-ended molecules) and sequenced to directly count template molecule abundance (Figure 1A). We use REcount to design a set of synthetic DNA standards that can be used to assess clustering bias due to molecule length on Illumina sequencers, and demonstrate that there is substantial variation in size bias between different Illumina instruments.

**Figure 1.**
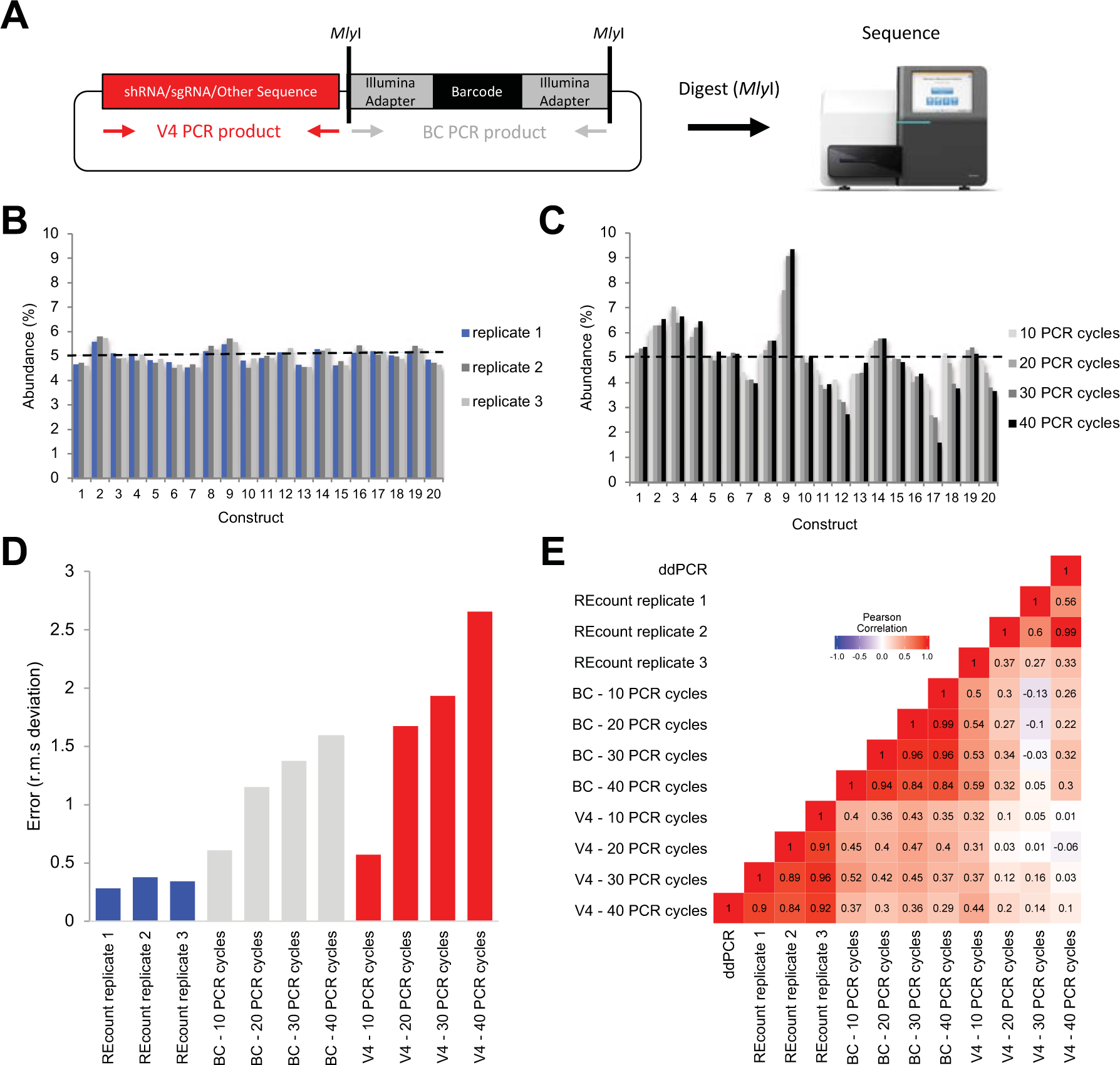
REcount enables accurate and precise measurements of plasmid pools. A) Design of REcount constructs. A barcode-containing, Illumina adapter-flanked construct is liberated with a restriction enzyme (*M/y*I) digest and directly sequenced. B) Accuracy and reproducibility of REcount. C) Analogous measurements of the same plasmid pool shown in panel B using varying PCR cycle numbers. D) Root mean squared deviation from expected values (5% per construct) when the plasmid pool is measured using REcount, and varying cycles of PCR amplification of either the barcode construct (BC) or another variable sequence in these plasmids (V4). E) Pearson correlation heatmap comparing REcount measurements with droplet digital PCR data and with conventional PCR amplification of either the BC or V4 amplicons.

## Results

In order to characterize the REcount method, we constructed a pool of 20 synthetic plasmids containing REcount barcodes, mixed at an equimolar abundance (5% per plasmid) based on fluorometric DNA concentration measurements. This pool was digested with *M/y*I and sequenced on an Illumina MiSeq. All 20 barcodes were detected at relative abundances ranging from 3.41% and 6.32% (CV = 0.13), consistent with the targeted abundances of 5% per construct (Supplemental Figure S1). To generate a more accurately pooled reference standard for subsequent experiments, we used this sequencing data as the basis for re-pooling the 20 plasmids and digested the new pool with *M/y*I and sequenced. The range of relative abundances of the re-pooled plasmids was narrower, ranging from 4.52% to 5.58% (CV = 0.06), indicating that the initial sequencing data was predictive in improving the accuracy of pooling as assessed by REcount (Supplemental Figure S1). To assess the reproducibility of these measurements, we digested and sequenced two additional replicates of the even plasmid pool. The replicate REcount measurements were highly reproducible with an average CV of 0.02 (Figure 1B).

Next, we compared REcount measurements of the even plasmid pool to PCR-based measurements, either of the barcode construct (BC) or another construct-specific sequence (V4). PCR-based measurements exhibited substantial construct-specific deviations from the expected 5% values, the extent of which increased with greater numbers of PCR cycles (Figure 1C-0). Furthermore, the construct-specific deviations from expected values were uncorrelated for the BC and V4 amplicon measurements, suggesting that the PCR biases were a function of template sequence (Supplemental Figure S2).

In order to independently measure the relative template concentrations in the even plasmid pool, we designed a pair of ddPCR assays targeting each barcode construct and validated the specificity of each assay using qPCR on each of the 20 individual plasmid templates (Supplemental Figure S3) (Hindson et al. 2011). The ddPCR-based measurements correlated well with the REcount measurements, both for the original and re-pooled even plasmid pools (Figure 1E, Supplemental Figure S3). In contrast, the PCR-based measurements of both the BC and V4 amplicons were not well-correlated with the ddPCR measurements (Figure 1E, Supplemental Figure S3). These results were corroborated with similar measurements of a pool of the same 20 plasmids mixed in a staggered manner, where PCR-based measurements had reduced correlation with ddPCR measurements and led to a systematic overestimation of the lower abundance constructs (Supplemental Figure S4). Taken together, these results indicate that REcount accurately reports on template abundance, while PCR-based measurements introduce increasing error with increased cycle numbers.

One drawback of the REcount method is that the indices that specify sample identity in multiplexed sequencing, which are typically flexibly added by PCR, are hard-coded into the constructs. To overcome this limitation, we tested whether orthogonal restriction enzymes could be used to multiplex REcount measurements. We initially chose *M/y*I as the flanking enzyme because it could precisely liberate the desired Illumina adapter-flanked construct. We tested whether other restriction enzymes that do not cleanly liberate flush Illumina adapter ends could also be used for REcount measurements. Initially, we tested *Bsm*I, *Bts^α^*I, and *Bsr*DI, each of which leave 2 nt 3’ overhangs. We constructed a pool of 12 plasmids comprised of sets of three barcoded constructs flanked by either *M/y*I, *Bsm*I, *Bts^α^*I, or *Bsr*DI (Figure 2A). In addition, all 12 of these constructs contained a pair of *Sbf*I sites located such that digestion with *Sbf*I liberates all 12 Illumina adapter-flanked cassettes with additional overhangs of between 30-36 bp upstream of the p5 flowcell adapter and between 40-50 bp downstream of the p? flowcell adapter. We digested this plasmid pool with each of the 5 enzymes individually and individually sequenced the digests and mapped the reads to a reference file containing all 12 expected barcodes. For *M/y*I, *Bsm*I, *Bts^α^*I, and *Bsr*DI, the expected barcodes were detected for each respective enzyme (Figure 2B-E, G-J). All 12 barcodes were detected when the pool was digested with *Sbf*I, indicating that clustering and sequencing can occur even in the presence of large (30-50 bp) overhangs (Figure F, K). We were not able to determine whether the length of the overhang affects the efficiency of clustering as each of these samples was sequenced in a portion of a MiSeq lane, together with other libraries. We observed differing amounts of off-target barcode detection in these orthogonal digests, ranging from <0.2% in the *Bsm*I digest to approximately 6% in the *Bts^α^*I digest (Figure 2B-E, G-J). This could likely be improved by adding a size selection step. Nonetheless, the fact that multiple restriction enzymes can be used to liberate REcount constructs allows for potential multiplexing strategies involving orthogonal digestion of distinct subpopulations of molecules or of concatamerized barcode arrays.

**Figure 2.**
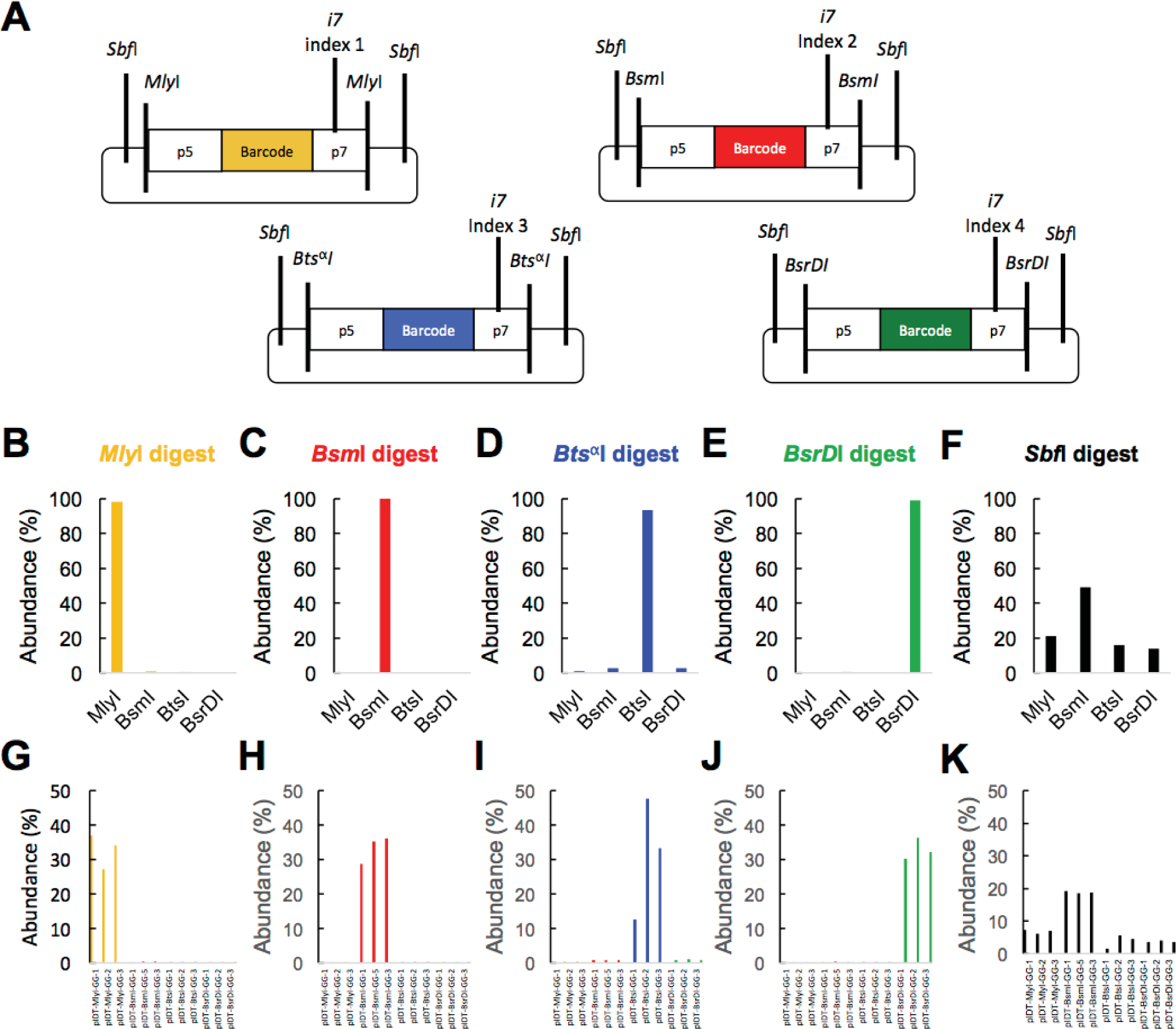
Multiplexing of REcount measurements using orthogonal restriction enzymes. A) Plasmids containing REcount constructs flanked by orthogonal restriction enzyme cut sites. B-F) Total mapped reads identified for each construct type when the plasmid pool is digested with the indicated enzyme. G-K) Mapped reads identified for each construct when the plasmid pool is digested with the indicated enzyme.

While it is known that molecule size affects clustering and sequencing efficiency on Illumina sequencers (Illumina 2014), the extent of this bias and the degree to which it differs between different Illumina instruments has not been characterized in detail. Thus, we used REcount to characterize the size bias profiles of the Illumina MiSeq, HiSeq 2500, HiSeq 4000, NextSeq, and NovaSeq sequencers. We synthesized 30 constructs, each of which contained an *M/y*I-flanked normalization barcode of consistent length (164 bp), and a barcode-containing variable-length insert ranging from 22 bp to 1372 bp, resulting in adapter-flanked molecules between 150 and 1500 bp (Figure 3A). In order to minimize sequence-specific artifacts, the variable-length inserts were chosen to have between 42% and 58% GC content, and were comprised of 10 constructs each (spanning the full 150 bp - 1500 bp size range) derived from three different molecules; the *Escherichia coli* (*E. coli*) 16S rRNA gene (16S), the *Drosophila melanogaster* (*D. melanogaster*) *alpha-Tubulin84B* gene (Tubulin), and the *D. melanogaster Glyceraldehyde-3-phosphate dehydrogenase 1* (GAPDH) gene (Supplemental Figure S5).

**Figure 3.**
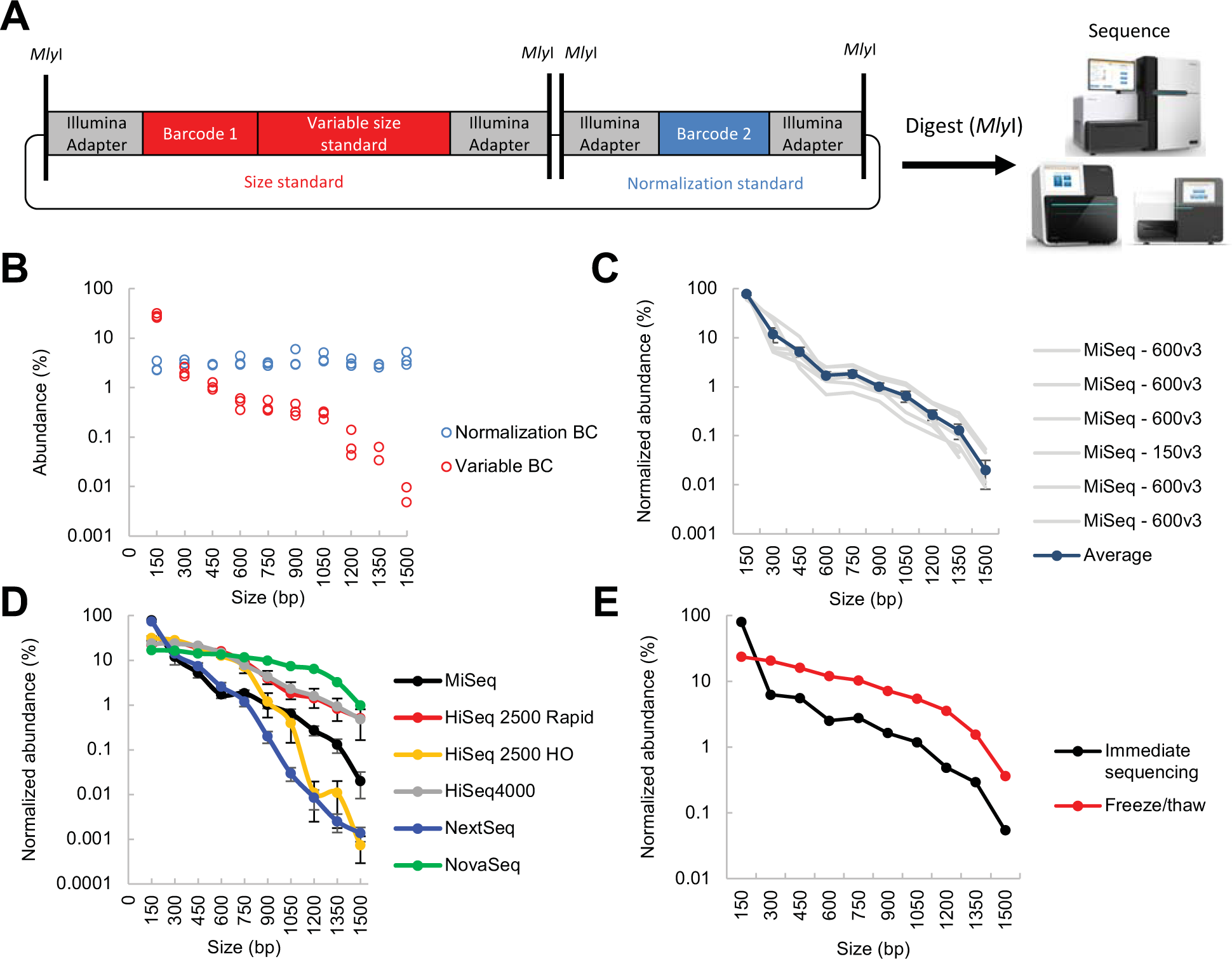
Illumina size standards allow measurement of sequencer-specific size biases. A) Design of REcount-based Illumina size standard constructs. Each standard construct contains a normalization barcode, as well as a barcode associated with a variable size standard that can be liberated by *M/y*I digestion and directly sequenced. B) Raw abundance data for all 30 size standards and normalization barcodes from a MiSeq run. C) Run-to-run variability of multiple MiSeq runs (n = 6 flow cells). D) Size bias profiles of the MiSeq (n=6 flow cells), HiSeq 2500 Rapid (n=1 flow cell, 2 lanes), HiSeq 2500 High Output (HO, n=2 flow cells, 10 lanes), HiSeq 4000 (n=3 flow cells, 6 lanes), NextSeq (n=4 flow cells), and NovaSeq (n=1 flow cell, 1 lane) sequencers. E) Size bias profiles of the same library either clustered on the MiSeq immediately after denaturation, or clustered after freezing and thawing the denatured library. Error bars are +/-s.e.m.

These Illumina size standard constructs were pooled at an equimolar ratio based on fluorometric DNA concentration measurements, digested with *M/y*I, and sequenced on different Illumina DNA sequencers with no intervening clean-up step, to ensure that no material was lost. Representative data from a single MiSeq run is shown in Figure 3B. Since each normalization barcode is present at an equimolar ratio to the corresponding size standard (as they are on the same plasmid), this allows any inaccuracies in plasmid pooling to be accounted for. Within a sequencing platform, clustering size bias exhibits run-to-run variation (Figure 3C). All five of the sequencers we tested exhibited preferential clustering of smaller fragments, consistent with previous anecdotal observations (Figure 3D). However, the magnitude of this effect and the shapes of the size bias curves differ substantially between the MiSeq, HiSeq 2500, HiSeq 4000, NextSeq, and NovaSeq (Figure 3D). Differences were also seen between the HiSeq 2500 in Rapid Run (onboard clustering) and High Output (cBot clustering) modes (Figure 3D). In addition, we observed an effect of molecule length on sequencing quality score, with a general trend towards longer molecules having lower quality scores (Supplemental Figure S5). The magnitude of the effect of molecule length on sequence quality varied among the different instruments.

The denaturation process can also affect the size bias observed on Illumina instruments. Denatured libraries are sometimes saved for re-sequencing in the case of a run failure (although Illumina’s best practices recommend preparing freshly denatured libraries). To test whether freshly denatured libraries perform differently from frozen denatured libraries, we sequenced a freshly denatured library on a MiSeq, and the same denatured library one day later, after a freeze-thaw cycle, on a second MiSeq. The freeze-thaw cycle had a substantial effect on the size bias profile of this library; in particular, there was a dramatic reduction in the fraction of 150 bp molecules observed, resulting in a corresponding upward shift of the curve (Figure 3E). It is likely that this shift reflects differential re-annealing of 150 bp fragments (which are in molar excess due to the presence of the large number of similarly sized normalization barcodes), or other small library molecules in the sequencing pool. This observation suggests that some of the difference in clustering size bias observed between the different platforms may be due to differences in denaturation conditions, the amount of time between loading the library and clustering, and whether the clustering process takes place in a chilled compartment (such as on the MiSeq) or not (such as the HiSeq2500 and NextSeq). Consistent with this idea, the variation between HiSeq2500 and HiSeq 4000 flow cells is much larger than the variation between the lanes on the same flow cell (Supplemental Figure S5, Supplemental Figure S6).

It is also likely that a portion of the variability between flow cells is due to differences in the size distributions of the libraries being sequenced together with the synthetic size standards, as competition for clustering will occur between all molecules in the sequencing lane. We observed a shift in the curve corresponding to a decreased representation of the larger size standards when they were sequenced together with a library containing a significant amount of material that was smaller than 300 bp on the HiSeq 4000 (Supplemental Figure 6). Although the size standards were sequenced together with different libraries across the different instruments, this context-dependent clustering is not sufficient to explain the large differences we see between different instruments. For example, libraries with similar average sizes and distributions yielded dramatically different measurements of size bias on the NextSeq versus the HiSeq 4000 (Supplemental Figure 6).

Surprisingly, we also detected an instance of construct-specific size bias, specifically on the HiSeq 2500 platform in Rapid Run mode (Supplemental Figure S5). In contrast to the MiSeq, HiSeq 2500 High Output, HiSeq4000, NextSeq, and NovaSeq where no systematic construct-specific biases were observed, the size bias curves for the 16S, GAPDH, and alpha-Tubulin constructs separated as size increased, with 16S showing much less of a drop-off with increased molecule size. One possible explanation for this difference is that the 16S rRNA gene has substantial secondary structure (Woese et al. 1980), which may serve to shorten the effective length of the molecule during the clustering process. This phenomenon may be due to differences in the clustering process or temperature on this platform, which may be less effective at dissociating the secondary structure of the 16S rRNA gene (https://support.illumina.com/bulletins/2016/10/considerations-when-migrating-nonillumina-libraries-between-sequencing-platforms.html). The HiSeq and MiSeq also have different recommended NaOH concentrations for denaturing libraries. It is possible that long molecules, particularly those with highly stable secondary structure, are incompletely denatured under the HiSeq denaturing conditions.

## Discussion

In summary, we describe REcount, a novel strategy for obtaining highly accurate and precise PCR-free NGS-based measurements of engineered genetic constructs. Similar constructs could be incorporated into shRNA, CRISPR, and transposon libraries to improve quantification of these molecules in pooled genetic screens. Currently, such measurements are prone to bias introduced by PCR, as we observed for both the BC and V4 amplicons (Figure 1, Supplemental Figure S4). Such sequence-specific amplification biases are often mitigated by including input controls, which are thought to accurately model amplification biases. However, amplification biases can be impacted by template concentration and by the context of the other molecules in the amplification reaction (Gohl et al. 2016), and can limit the sensitivity of these assays by compressing the dynamic range (Supplemental Figure S4). One challenge of employing PCR-free quantification barcodes in these contexts is the large amount of genomic DNA relative to the PCR-free barcode construct. However, we have shown that transposon pools can be quantified from isolated *E. coli* genomic DNA using this approach (data not shown). We further demonstrated that multiplexing of REcount measurements is possible using orthogonal restriction enzymes (Figure 2).

We used REcount to measure size bias on several different Illumina sequencers. We found that size bias can vary between runs and instruments and that the denaturation procedure can affect the size bias (Figure 3). Due to the competitive clustering of molecules of different sizes, it is likely that a portion of the variability between runs and lanes is due to differences in the size distributions of the libraries being sequenced together with the synthetic size standards. Thus, the shape of the size bias curve is likely sensitive to both the size distribution of the libraries being sequenced along with the size standards, as well as the proportion of the lane devoted to the size standards.

While such size biases would not be expected to affect randomly sheared libraries, these results indicate that care should be taken when interpreting quantitative measurements or comparing data across different platforms. This is particularly true in cases where library size distributions are non-random such as in several chromatin profiling methods (e.g. ATAC-Seq (Buenrostro et al. 2013), FAIRE-Seq/MAINE-Seq (Ponts et al. 2010)), approaches that use restriction digestion to fragment DNA (e.g. RAD-Seq (Andrews et al. 2016)), amplicons that vary in length (e.g. fungal ITS sequencing (Taylor et al. 2016)), or techniques such as TAIL-Seq (Chang et al. 2014) that explicitly seek to measure molecule length. Constructs such as those described here could be routinely spiked into Illumina sequencing runs to monitor size bias, similar to the use of PhiX to report on sequencing error rates and other base-calling metrics.

## Methods

### Synthesis and cloning of REcount plasmids

#### Even and staggered pool plasmids

The plasmids comprising the even and staggered pools were designed to include a portion of the 16S rRNA gene from one of twenty different bacterial species, modeled on the Human Microbiome Project mock microbial communities (HM-276D and HM-277D, (2012a, 2012b), with a 3 bp “TCT” sequence tag added at an analogous position in each construct. These constructs also contained an I-SceI site, allowing for linearization of the plasmids, and a REcount construct, consisting of a unique 20 bp DNA barcode, flanked by Illumina adapters and *M/y*I restriction sites, spaced in a manner to precisely liberate the Illumina adapter-containing barcode construct (Supplemental_File_1). These constructs were synthesized as DNA tiles by SGI-DNA and assembled into full-length constructs using the BioXP 3200 (SGI-DNA). The assembled DNA fragments were A-tailed using the A-tailing module from NEB, cloned into pCR2.1 using a TOPO TA cloning kit (Thermo Scientific), and transformed into OneShot TOP10 chemically competent *E. co/i* (Thermo Scientific). Multiple colonies were selected, DNA was isolated using a Qiagen MiniPrep kit, and sequence-verified clones were identified by Sanger sequencing with the following primers:

M13F: GTAAAACGACGGCCAG

M13R: CAGGAAACAGCTATGAC

The twenty sequence-verified plasmids were quantified using a Quant-iT PicoGreen dsDNA assay (Thermo Fisher Scientific), normalized to 50 ng/µl, and pooled at equal volume to create the original even pool. The re-pooled even pool and staggered pool were made by adjusting the volume pooled based on the initial PCR-free sequencing data of the original even pool.

#### Orthogonal enzyme multiplexing plasmids

Four synthetic gene fragments were synthesized (IDT) in the pIDTSmart-Amp plasmid backbone, consisting of an Illumina adapter containing construct with internal *Pac*I and *Pme*I site, and flanked by a pair of either *M/y*I, *Bsm*I, *Bts*^α^I, or *Bsr*DI sites. The full constructs were also flanked by a pair of *Sbf*I sites (Supplemental File 2). In order to make a collection of barcode-containing constructs, the plasmid templates were amplified using the following template-specific primers, and a Golden Gate cloning reaction was used to re-generate the circular plasmid:

UMGC_350_MlyI_barcode_p5:

NNNNGGTCTCTACTTATCCWWNNNWWNNNAGATCGGAAGAGCGTCGTGTAG

UMGC_350_MlyI_barcode_p7:

NNNNGGTCTCTAAGTGCAANNNWWNNNWWAGATCGGAAGAGCACACGTCTGAA

UMGC_350_BsmI_barcode_p5:

NNNNGGTCTCTGGTTATCCNNSSNNSSNNAGATCGGAAGAGCGTCGTGTAG

UMGC_350_BsmI_barcode_p7:

NNNNGGTCTCTCCAAGCAANNSSNNSSNNAGATCGGAAGAGCACACGTCTGAA

UMGC_350_BtsI_barcode_p5:

NNNNGGTCTCTGAACATCCNNNWWNNNWWAGATCGGAAGAGCGTCGTGTAG

UMGC_350_BtsI_barcode_p7:

NNNNGGTCTCTGTTCGCAANNNWWNNNWWAGATCGGAAGAGCACACGTCTGAA

UMGC_350_BsrDI_barcode_p5:

NNNNGGTCTCTATGAATCCNNSSNNSSNNAGATCGGAAGAGCGTCGTGTAG

UMGC_350_BsrDI_barcode_p7:

NNNNGGTCTCTTCATGCAANNSSNNSSNNAGATCGGAAGAGCACACGTCTGAA

Briefly, PCR reactions were set up using the following recipe: 1 µl plasmid DNA (20 ng/µl), 2.5 µl primer 1 (10 µM), 2.5 µl primer 2 (10 µM), 19 µl water, and 25 µl 2x Q5 master mix (NEB). PCR amplification was carried out using the following cycling conditions: 98**°**C for 30 seconds, followed by 30 cycles of 98**°**C for 20 seconds, 60**°**C for 15 seconds, 72**°**C for 1.5 minutes, followed by 72**°**C for 5 minutes. Golden Gate reactions(Engler et al. 2008, 2009) were set up using the following recipe:

1 µl Barcoding PCR product from above, 2 µl NEB Cutsmart buffer, 2 µl 10mM ATP (NEB), 12.5 µl nuclease-free water, 0.5 µl *Bsa*I-HF, 1 µl T4 DNA ligase (NEB 400,000 U/ml), 1 µl *Pac*I. Golden Gate reactions were cycled with the following conditions: 10 cycles of 37**°**C for 5 minutes, 21**°**C for 5 minutes, then 1 cycle 37**°**C for 10 minutes, then 1 cycle 80**°**C for 20 minutes. Golden Gate reactions were transformed into chemically competent *E. coli* 5alpha cells (NEB). Colonies were picked and DNA was isolated using a Qiagen Mini-Prep kit. Uniquely barcoded constructs were identified by Sanger sequencing with the following primers:

UMGC_350-pIDT-Smart-For: CTGAGGCTCGTCCTGAATGATA

UMGC_350-pIDT-Smart-Rev: ACCGATCATACGTATAATGCCGTAA

The twelve sequence-verified plasmids were quantified using a Quant-iT PicoGreen dsDNA assay (Thermo Fisher Scientific), normalized to 50 ng/µl, and pooled at equal volume to create the orthogonal enzyme multiplexing test pool. Subsequent NGS analysis indicated that some of these clones were mixed isolates, as other barcodes that had not been detected by Sanger sequencing were present in the NGS data sets. Analysis is based on the Sanger-verified barcodes only.

#### Illumina size standard plasmids

Illumina size standards were designed using three different template molecules as backbones for the variable length fragment; the 16S rRNA gene (16S) from *E. coli*, the *alpha-Tubulin84B* gene (Tubulin) from *D. melanogaster*, and the *Glyceraldehyde-3-phosphate dehydrogenase 1* (GAPDH) gene from *D. melanogaster* (Supplemental Figure S5). Any naturally occurring *M/y*I sites in these fragments were modified to remove this restriction site. The variable length size standards represent nested fragments of these three genes with breakpoints chosen to generate specific molecule lengths, with GC contents between 40-60% (Figure 3, Supplemental Figure S5). In order to minimize repetitive sequences, different adapters were used for the normalization and variable size standards (Nextera and TruSeq, respectively), and the normalization and size standards were synthesized in opposite orientations in the construct. Both the Illumina adapter flanked variable and normalization barcode constructs were flanked by *M/y*I restriction sites. The Illumina size standard constructs were synthesized by GenScript in the pUC57 cloning vector (Supplemental File 3). Approximately 4 µg of each lyophilized plasmid was resuspended in 40 µl of EB (Qiagen). Plasmids were quantified using a Quant-iT PicoGreen dsDNA assay (Thermo Fisher Scientific) and normalized to 10 nM to account for the variable sizes of the plasmids, then pooled at an equimolar ratio.

### qPCR validation of ddPCR assays

A set of primers allowing amplification between the construct-specific barcode and the Illumina flow cell adapter, either in the forward orientation (assay 1, where the construct-specific primer was paired with the p7 primer) or reverse orientation (assay 2, where the construct-specific primer was paired with the p5 primer) were designed and synthesized (Integrated DNA Technologies, Supplemental File 4). In order to validate these assays, we performed qPCR amplification of each individual plasmid, the even plasmid pool, and a negative control (water) with each of the 40 primer sets, as well as a p5/p7 positive control (which is expected to amplify all constructs). PCR reactions were set up as follows: 3 µl template DNA (0.05 ng/µl), 1.06 µl nuclease-free water, 0.6 µl 10x Qiagen PCR buffer, 0.24 µl MgCl_2_ (25 mM), 0.3 µl DMSO, 0.048 µl dNTPs (25 mM), 0.12 µl ROX (25 µM), 0.003 µl SYBR (1000x), 0.03 µl Qiagen Taq (5 U/µl), 0.3 µl primer 1 (10 µM), and 0.3 µl primer 2 (10 µM). Reactions were amplified on an ABI 7900 with the following cycling conditions: 95ºC for 5 minutes, followed by 35 cycles of: 94ºC for 30 seconds, 55ºC for 30 seconds, and 72ºC for 30 seconds, followed by incubation at 72ºC for 1 minute. For each primer set, Ct values were normalized to the mean Ct for that primer set across all plasmids and plotted as a heatmap (Supplemental Figure S5).

### ddPCR

The re-pooled even plasmid mix was quantified using a Quant-iT PicoGreen dsDNA assay (Thermo Fisher Scientific), diluted to 1 ng/µl, and further diluted 1:10,000 to bring the pool to the correct concentration for digital quantification. The following ddPCR reactions were prepared: 5 µl template DNA, 0.44 µl primer 1 (10 µM), 0.44 µl primer 2 (10 µM), 5.12 µl water, and 11 µl EvaGreen reaction mix (BIO-RAD). In addition, 2 µl of */-See/* was added to the ddPCR master mix to linearize the plasmid DNA templates, resulting in between 0.02 and 0.075 µl of */-See/* per reaction. Emulsion droplets were generated using a QX200 Droplet Generator (BIO-RAD) following the manufacturer’s instructions, transferred to a 96-well PCR plate, and cycled using the following conditions: 95ºC for 10 minutes, followed by 40 cycles of: 95ºC for 30 seconds and 55ºC for 1 minute, followed by a final extension step of 72ºC for 5 minutes, and a 12ºC hold.

Droplets were counted using a QX200 Droplet Reader (BIO-RAD). The re-pooled even plasmid mix was run in triplicate for both the forward and reverse assays. Single replicates of both the original even pool and the staggered pool were run for both assays. For the staggered pool, the extent of dilution of the 1 ng/µl plasmid pool was varied such that the template abundance of the plasmid targeted by the primer set was expected to be at the correct concentration for digital quantification. Data was analyzed using QuantaSoft Analysis Pro software (BIO-RAD). Replicate measurements were averaged (when available) for both ddPCR assays in order to arrive at a measurement of average ddPCR counts for each construct. Data from the assay was not included in cases where there was not clear separation between positive and negative droplets.

### Sequencing library preparation

#### Even and staggered pool REcount measurements

The following *M/y*I digests were set up for PCR-free quantification: 200-500 ng even or staggered pool DNA, 2 µl Cutsmart buffer (NEB), 1 µl MlyI (NEB), and volume was adjusted to 20 µl with nuclease-free water. Digests were incubated at 37ºC for 1 hour, followed by 20 minutes at 65ºC. 30 µl of water was added to each digest (to bring the volume up to 50 µl). 30 µl (0.6x) of AmpureXP beads (Beckman Coulter) were added and after a 5 minute incubation, beads were collected on a magnet and the supernatant was transferred to a new tube (discarded beads). 80 µl (1x) of AmpureXP beads was added, washed 2x for 30 seconds using fresh 80% ethanol, and beads were air dried for 10 minutes, followed by elution in 20 µl of EB (Qiagen). Libraries were quantified using a Quant-iT PicoGreen dsDNA assay (Thermo Fisher Scientific), fragment sizes were assessed using an Agilent Bioanalyzer High Sensitivity assay, and libraries were normalized to 2 nM for sequencing.

#### Even and staggered pool PCR-based measurements

##### Barcode construct (BC) library preparation

The following PCR reactions were set up to amplify the BC constructs: 1 µl DNA (1 ng/µl), 5 µl 10x Qiagen PCR buffer, 2 µl MgCl_2_ (25 mM), 2.5 µl DMSO, 0.4 µl dNTPs (25 mM), 0.25 µl Qiagen Taq (5 U/µl), 2.5 µl primer 1 (10 µM), 2.5 µl primer 2 (10 µM), and 33.85 µl nuclease-free water.

The following primers were used to amplify the BC constructs:
p5: AATGATACGGCGACCACCGA
p7: CAAGCAGAAGACGGCATACGA

Samples were amplified using the following cycling conditions: 95ºC for 5 minutes, followed by 10, 20, 30, or 40 cycles of 94ºC for 30 seconds, 55ºC for 30 seconds, and 72ºC for 30 seconds, followed by incubation at 72ºC for 10 minutes. Libraries were quantified using a Quant-iT PicoGreen dsDNA assay (Thermo Fisher Scientific), fragment sizes were assessed using an Agilent Bioanalyzer High Sensitivity assay, and libraries were normalized to 2 nM for sequencing.

##### V4 fragment library preparation

The following PCR reactions were set up in triplicate to amplify the V4 constructs: 2 µl DNA (0.1 ng/µl), 0.5 µl primer 1 (10 µM), 0.5 µl primer 2 (10 µM), 2 µl nuclease-free water, and 5 µl 2x Q5 master mix. The following primers were used:

V4_515F_Nextera:

TCGTCGGCAGCGTCAGATGTGTATAAGAGACAGGTGCCAGCMGCCGCGGTAA

V4_806R_Nextera:

GTCTCGTGGGCTCGGAGATGTGTATAAGAGACAGGGACTACHVGGGTWTCTAAT

Reactions were amplified using the following cycling conditions: 98**°**C for 30 seconds, followed by 10, 20, 30, or 40 cycles of 98**°**C for 20 seconds, 55**°**C for 15 seconds, 72**°**C for 1 minute, followed by 72**°**C for 5 minutes.

After initial amplification, PCR reactions were diluted 1:60 in nuclease-free water, and used as templates in the following indexing reactions: 3 µl PCR 1 (1:60 dilution), 1 µl indexing primer 1 (5 µM), 1 µl indexing primer 2 (5 µM), and 5 µl 2x Q5 master mix. The following indexing primers were used (X indicates the positions of the 8 bp indices):

Forward indexing primer:

AATGATACGGCGACCACCGAGATCTACACXXXXXXXXTCGTCGGCAGCGTC

Reverse indexing primer:

CAAGCAGAAGACGGCATACGAGATXXXXXXXXGTCTCGTGGGCTCGG

Reactions were amplified using the following cycling conditions: 98**°**C for 30 seconds, followed by 10 cycles of 98**°**C for 20 seconds, 55**°**C for 15 seconds, 72**°**C for 1 minute, followed by 72**°**C for 5 minutes. The full indexing PCR reactions were then purified and normalized using a SequalPrep normalization plate (Thermo Fisher Scientific), followed by elution in 20 µl of elution buffer. An even volume of the normalized libraries was pooled and concentrated using 1x AmpureXP beads (Beckman Coulter). Pooled libraries were quantified using a Qubit dsDNA broad-range assay (Thermo Fisher Scientific), fragment sizes were assessed using an Agilent Bioanalyzer High Sensitivity assay, and libraries were normalized to 2 nM for sequencing.

#### Orthogonal enzyme multiplexing tests

The twelve-plasmid orthogonal enzyme pool was cut with one of 5 different enzymes (in separate reactions) using the following recipe and enzyme-specific incubation conditions: 20 µl DNA (1 µg), 4 µl NEB buffer (CutSmart or NEB 2.1, depending on enzyme), 2 µl Enzyme (either *M/y*I [37**°**C for 1 hour, followed by 65**°**C for 20 minutes], *Bsm*I [65**°**C for 1 hour, followed by 80**°**C for 20 minutes], *Bts*^α^I [55**°**C for 1 hour], or *Bsr*DI [65**°**C for 1 hour, followed by 80**°**C for 20 minutes], or *Sbf*I [37**°**C for 1 hour, followed by 80**°**C for 20 minutes]), and 14 µl water. 10 µl (0.5x) of AmpureXP beads (Beckman Coulter) was added to 20 µl of digested DNA and after a five minute incubation, the beads were collected on a magnet and the supernatant was transferred to new tube (discarded beads). 20 µl of AmpureXP beads was added, and the beads were washed 2x for 30 seconds using fresh 80% ethanol, then air dried for 10 minutes, before eluting in 20 µl of EB (Qiagen). Libraries were quantified using a Quant-iT PicoGreen dsDNA assay (Thermo Fisher Scientific), fragment sizes were assessed using an Agilent Bioanalyzer High Sensitivity assay, and libraries were normalized to 2 nM for sequencing.

#### Illumina size standards

The following digest of the Illumina size standard pool was set up: 175 µl DNA (10 nM), 20 µl CutSmart buffer (NEB), 5 µl *M/y*I (NEB). The reaction was incubated at 37**°**C for 1 hour, followed by 65**°**C for 20 minutes. The library was quantified using a Quant-iT PicoGreen dsDNA assay (Thermo Fisher Scientific), fragment sizes were assessed using an Agilent Bioanalyzer High Sensitivity assay, and libraries were normalized to 2 nM for sequencing.

### Sequencing

DNA libraries were denatured with NaOH and prepared for sequencing according to the protocols described in the Illumina MiSeq, NextSeq, HiSeq 2500, HiSeq 4000, and NovaSeq Denature and Dilute Libraries Guides. Libraries were generally sequenced along with other samples in a fraction of a sequencing lane.

### Data analysis

REcount data was analyzed using custom R and Python scripts and BioPython (Cock et al. 2009). The first 20 bp of the sequencing reads was mapped against a barcode reference file (Supplemental Files 5-8), with a maximum of 2 mismatches allowed. A generalized script for counting REcount barcodes contained in a reference file is available on Github (https://github.com/darylgohl/REcount). Analysis of the V4 amplicon data was performed using the reference-based mapping pipeline described here: https://bitbucket.org/jgarbe/gopher-pipelines/wiki/metagenomics-pipeline.rst, using the reference file in Supplemental File 9 to build the bowtie2 index (Langmead and Salzberg 2012). For the analysis of quality scores (Supplemental Figure S5), the data for all runs on a given platform was concatenated into a single fastq file, the split into individual fastq files for each individual construct, based on the 20 bp sequence barcodes in each construct. Next, the reads were trimmed to 50 bp using cutadapt (Martin 2011), so that all constructs and sequencing runs could be compared in a standardized manner. Mean quality scores were calculated for each construct that was represented by at least 100 reads in the data set.

### Data access

Sequencing data files are available through the NCBI Sequence Read Archive (BioProject: PRJNA431017).

## Acknowledgements

We thank the staff of the University of Minnesota Genomics Center for advice, technical support, and data generation, and the Vincent J. Coates Genomics Sequencing Laboratory at UC Berkeley for data generation. We also thank Alessandro Magli, Daniel Schmidt, Igor Libourel, Steve Bowden, Benjamin Auch, John Garbe, and Nagendra Palani for helpful discussions. This work was supported by a grant from the University of Minnesota-Mayo Translational Product Development Fund to D.M.G. and K.B.B. (National Center for Advancing Translational Sciences of the National Institutes of Health Award Number UL1TR000114). The Vincent J. Coates Genomics Sequencing Laboratory at UC Berkeley was supported by an NIH S10 OD018174 Instrumentation Grant.

## Author contributions

D.M.G. and K.B.B. conceived and designed the experiments, analyzed data, and wrote the manuscript. D.M.G., A.B., D.J., S.A., B.B., and S.M. conducted the experiments.

## Disclosure declaration

The REcount PCR-free quantification barcode technology described here is included in US patent application numbers 62/332,879, 62/630,463, and PCT/US17/31271. D.M.G. is a shareholder and CSO of CoreBiome, Inc. K.B.B. is a shareholder and COO of CoreBiome, Inc.

## Supplemental Data Files for

**Measuring Illumina Size Bias Using REcount: A Novel Method for Highly Accurate Quantification of Engineered Genetic Constructs**

I. Supplemental Figures
  Supplemental Figure 1 Initial and re-pooled even plasmid pool data.
  Supplemental Figure 2 Lack of correlation between BC and V4 PCR.
  Supplemental Figure 3 Droplet digital PCR assay validation and data.
  Supplemental Figure 4 Assessment of REcount measurements of a staggered plasmid pool.
  Supplemental Figure 5 Illumina size standard pool composition and data.
  Supplemental Figure 6 Context-specific effects on clustering of size standards.
II. Supplemental Files
  1. Supplemental_File_1.fasta Sequences of the synthetic DNA standards used to construct the even and staggered plasmid pools.
  2. Supplemental_File_2.fasta Sequences of the synthetic DNA used to construct the orthogonal restriction enzyme plasmids.
  3. Supplemental_File_3.fasta Sequences of the synthetic DNA inserts from the Illumina size standard plasmids.
  4. Supplemental_File_4.xlsx A table of the primers used for the ddPCR assays in this study.
  5. Supplemental_File_5.fasta REcount barcode mapping file from the even and staggered plasmid pools.
  6. Supplemental_File_6.fasta Expected REcount barcodes for orthogonal enzyme multiplexing tests.
  7. Supplemental_File_7.fasta Normalization barcode mapping file for Illumina size standards.
  8. Supplemental_File_8.fasta Variable barcode mapping file for Illumina size standards.
  9. Supplemental_File_9.fasta Reference sequences for V4 PCR mapping.

**Supplemental Figure 1.**
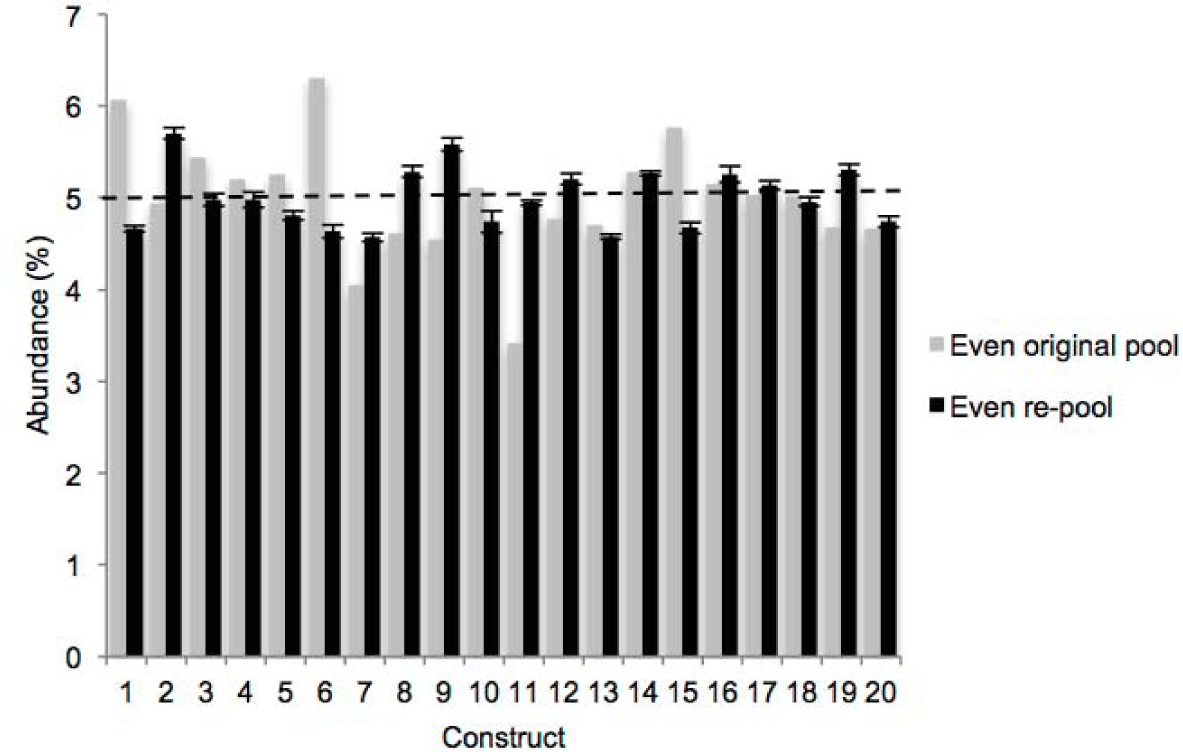
Initial and re-pooled even plasmid pool data. REcount measurements of an initial attempt at even plasmid pooling based on PicoGreen data, and a subsequent re-pooling informed by the initial pool sequencing data.

**Supplemental Figure 2.**
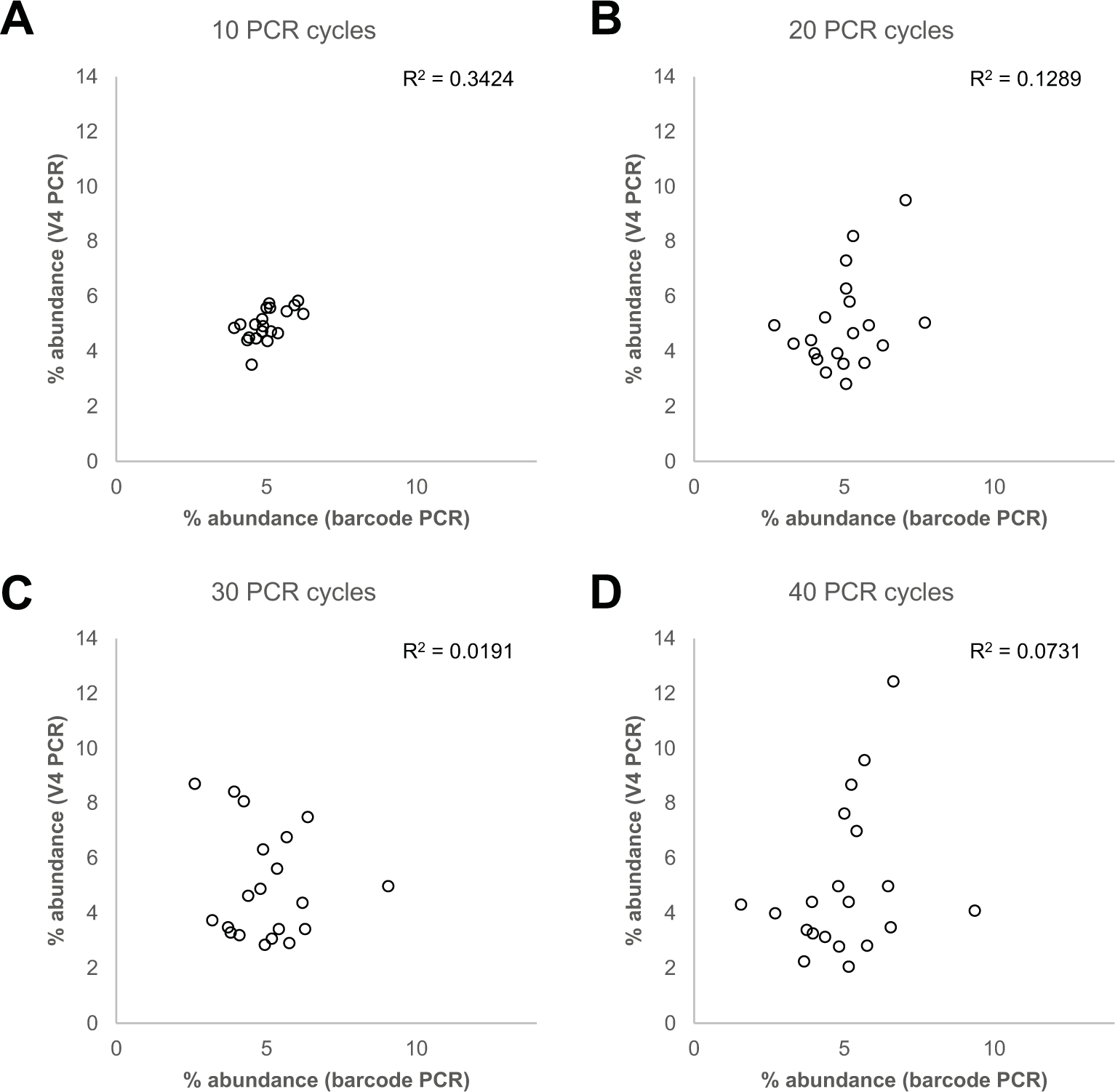
Lack of correlation between BC and V4 PCR. Scatterplots of BC and V4 abundance data for the even plasmid pool, when amplified for A) 10, B) 20, C) 30, or D) 40 PCR cycles.

**Supplemental Figure 3.**
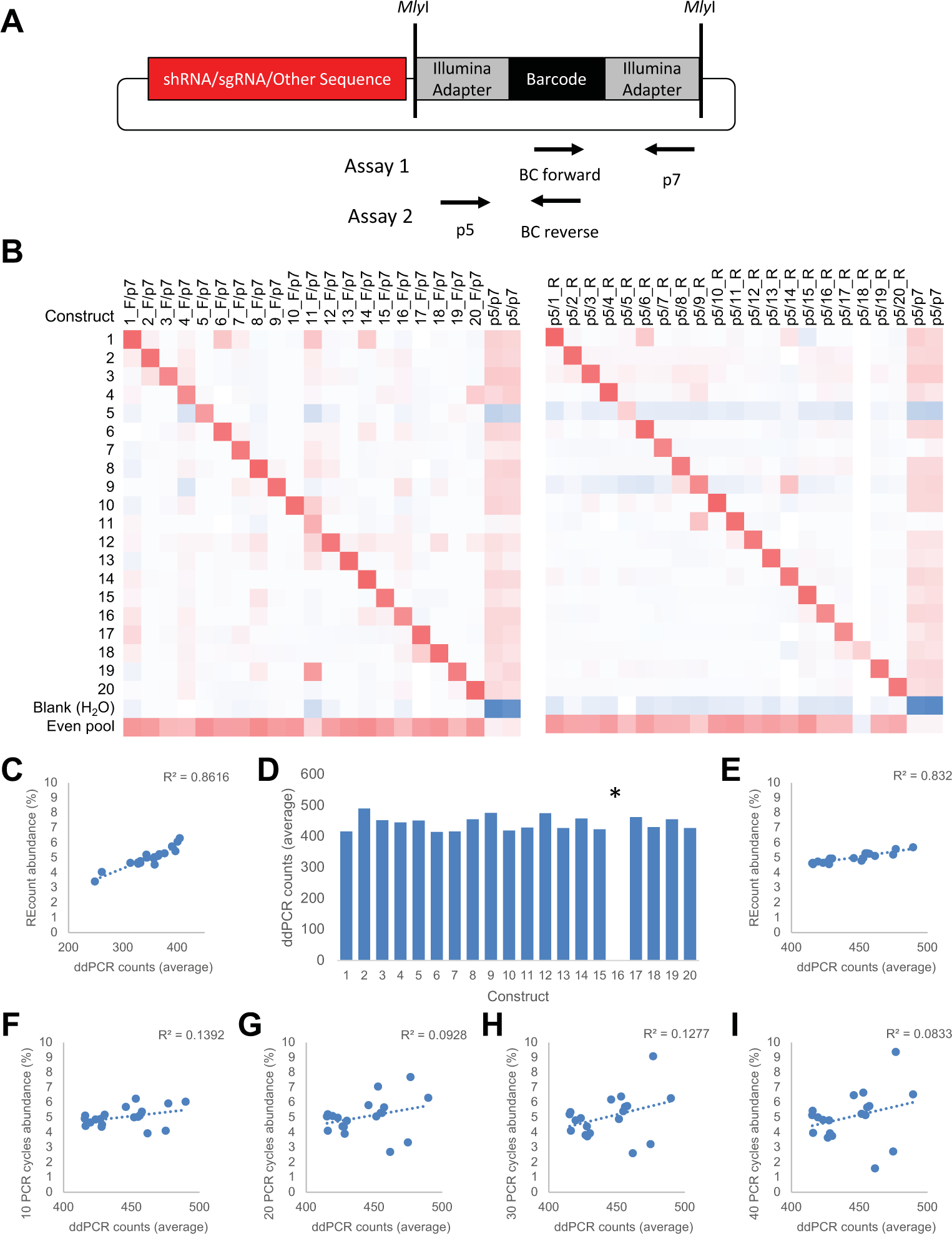
Droplet digital PCR assay validation and data. A) Schematic depicting the two ddPCR assays that were developed for each construct in the plasmid pool. B) qPCR data showing the specificity of each assay for the target construct as assessed by amplification of each individual plasmid, the even plasmid pool, or a negative control, with each primer pair. C) Correlation between ddPCR data and REcount quantification for the original even plasmid pool. D) ddPCR counts for the re-pooled even plasmid pool. Bars are the average of triplicates of the forward and reverse ddPCR assays where data could be generated for both assays, or just the forward or reverse assay in the case where one assay failed. *For plasmid 16, both the forward and reverse assays failed and thus no ddPCR information is available for this construct. E) Correlation between ddPCR data and REcount quantification for the re-pooled even plasmid pool. F-I) Correlation between ddPCR data and BC PCR-based quantification of the re-pooled even plasmid pool amplified for F) 10, G) 20, H) 30, I) 40 PCR cycles.

**Supplemental Figure 4.**
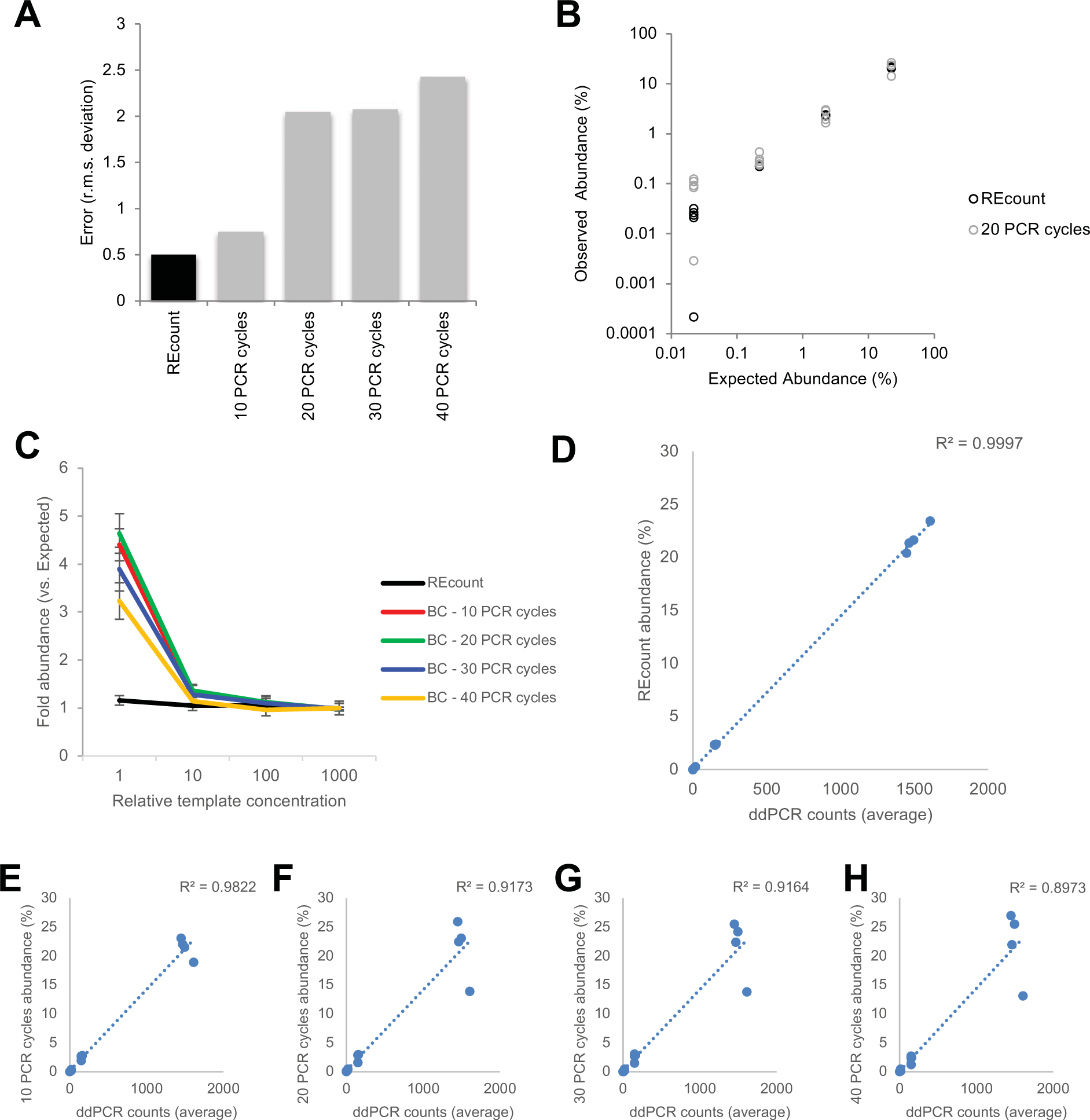
Assessment of REcount measurements of a staggered plasmid pool. A) Root mean squared deviation from expected values when the staggered plasmid pool is measured using REcount, and varying cycles of PCR amplification of the barcode construct. B) Comparison of REcount and PCR-based measurements of the staggered plasmid pool. C) Average measured representation of constructs pooled at different levels relative to expected values when measured using REcount or varying cycles of PCR. D) Correlation of ddPCR data and REcount measurements of the staggered plasmid pool. E-H) Correlation between ddPCR data and BC PCR-based quantification of the staggered plasmid pool amplified for E) 10, E) 20, G) 30, H) 40 PCR cycles. Error bars are +/-s.e.m.

**Supplemental Figure 5.**
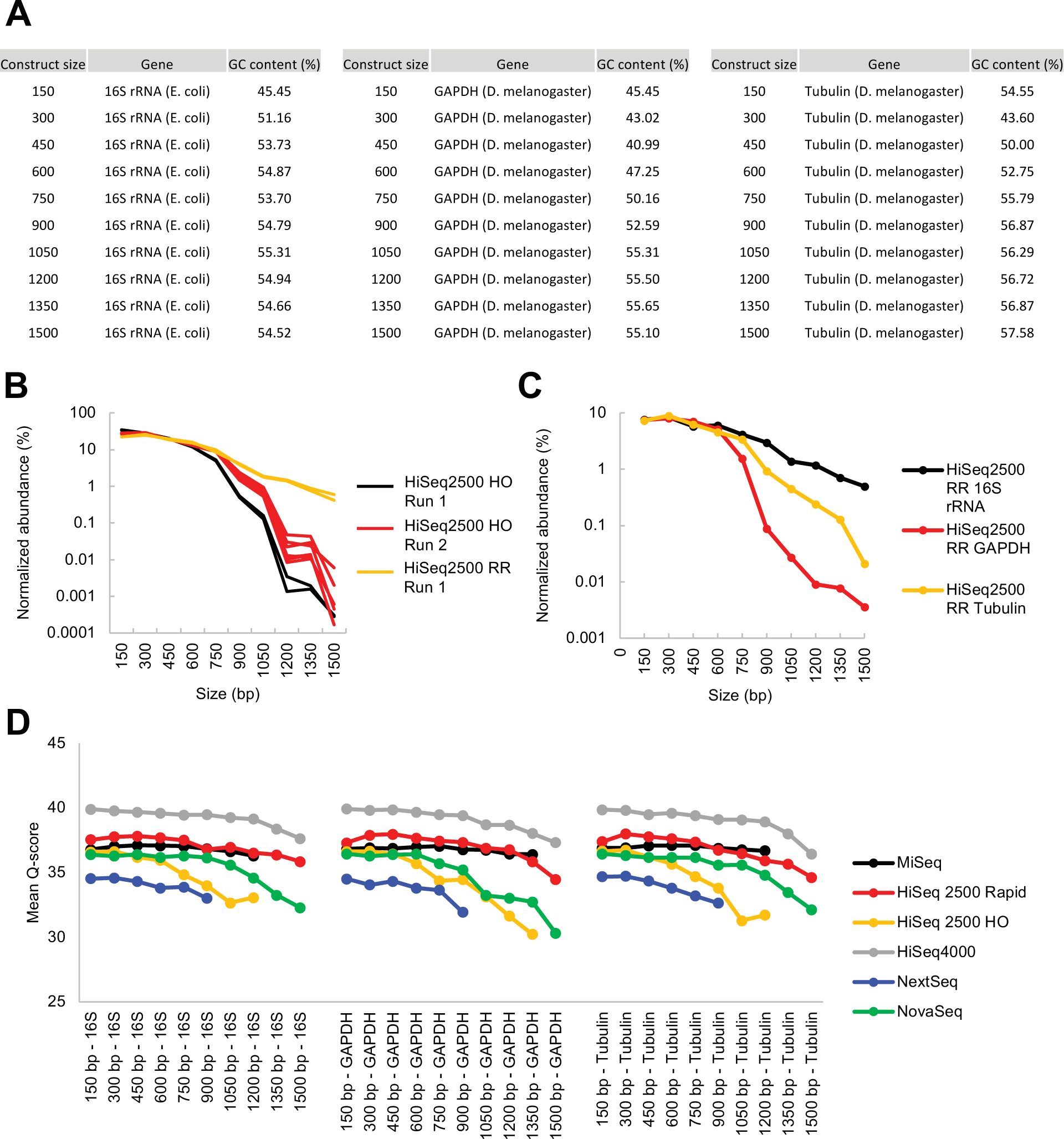
Illumina size standard pool composition and data. A) Composition of the Illumina size standard constructs, which consist of three different backbone molecules (16S rRNA, GAPDH, and Tubulin), ranging from 150 bp to 1500 bp in length. B) Between lane and between flow cell differences in size bias profiles for HiSeq2500 Rapid Run (on-board clustering) and HiSeq2500 High Output (cBot clustering). C) Template-specific size biases observed on the HiSeq2500 in Rapid Run mode. D) Platform and construct-specific mean quality scores for the Illumina size standard constructs for the first 50 bp of read 1.

**Supplemental Figure 6.**
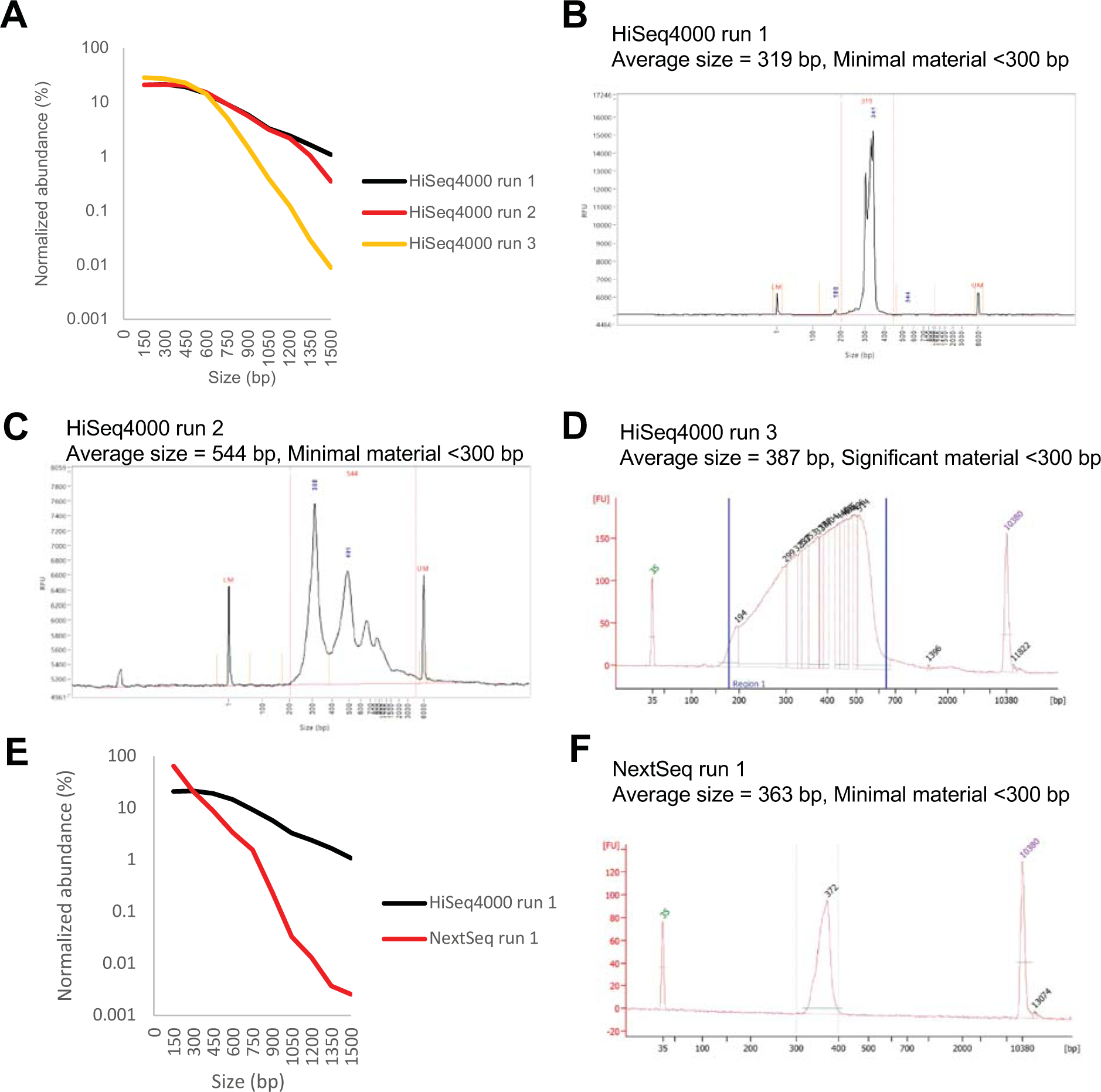
Context-specific effects on clustering of size standards. A) Differences in size standard measurements for three HiSeq 4000 runs (3 different flow cells). B) Fragment size profile of the library run together with the size standards in run 1 of the HiSeq 4000. C) Fragment size profile of the library run together with the size standards in run 2 of the HiSeq 4000. D) Fragment size profile of the library run together with the size standards in run 3 of the HiSeq 4000. E) Differences in size standard measurements for run 1 of the HiSeq 4000 and run 1 of the NextSeq. F) Fragment size profile of the library run together with the size standards in run 1 of the NextSeq.

## References

1. Aird D, Ross MG, Chen W-S, Danielsson M, Fennell T, Russ C, Jaffe DB, Nusbaum C, Gnirke A. 2011. Analyzing and minimizing PCR amplification bias in Illumina sequencing libraries. Genome Biol 12: R18.

2. Andrews KR, Good JM, Miller MR, Luikart G, Hohenlohe PA. 2016. Harnessing the power of RADseq for ecological and evolutionary genomics. Nat Rev Genet 17: 81–92.

3. Bhang HC, Ruddy DA, Krishnamurthy Radhakrishna V, Caushi JX, Zhao R, Hims MM, Singh AP, Kao I, Rakiec D, Shaw P, et al. 2015. Studying clonal dynamics in response to cancer therapy using high-complexity barcoding. Nat Med 21: 440–448.

4. Buenrostro JD, Giresi PG, Zaba LC, Chang HY, Greenleaf WJ. 2013. Transposition of native chromatin for fast and sensitive epigenomic profiling of open chromatin, DNA-binding proteins and nucleosome position. Nat Methods 10: 1213–1218.

5. Chang H, Lim J, Ha M, Kim VN. 2014. TAIL-seq: genome-wide determination of poly(A) tail length and 3’ end modifications. Mol Cell 53: 1044–52.

6. Cock PJA, Antao T, Chang JT, Chapman BA, Cox CJ, Dalke A, Friedberg I, Hamelryck T, Kauff F, Wilczynski B, et al. 2009. Biopython: freely available Python tools for computational molecular biology and bioinformatics. Bioinformatics 25: 1422–3.

7. Engler C, Gruetzner R, Kandzia R, Marillonnet S. 2009. Golden Gate Shuffling: A One-Pot DNA Shuffling Method Based on Type IIs Restriction Enzymes ed. J. Peccoud. PLoS One 4: e5553.

8. Engler C, Kandzia R, Marillonnet S. 2008. A one pot, one step, precision cloning method with high throughput capability. ed. H.A. El-Shemy. PLoS One. 3: e3647.

9. Geiss GK, Bumgarner RE, Birditt B, Dahl T, Dowidar N, Dunaway DL, Fell HP, Ferree S, George RD, Grogan T, et al. 2008. Direct multiplexed measurement of gene expression with color-coded probe pairs. Nat Biotechnol 26: 317–325.

10. Gohl DM, Vangay P, Garbe J, MacLean A, Hauge A, Becker A, Gould TJ, Clayton JB, Johnson TJ, Hunter R, et al. 2016. Systematic improvement of amplicon marker gene methods for increased accuracy in microbiome studies. Nat Biotechnol 34: 942–949.

11. Hindson BJ, Ness KD, Masquelier DA, Belgrader P, Heredia NJ, Makarewicz AJ, Bright IJ, Lucero MY, Hiddessen AL, Legler TC, et al. 2011. High-Throughput Droplet Digital PCR System for Absolute Quantitation of DNA Copy Number. Anal Chem 83: 8604–8610.

12. Illumina. 2014. Nextera^®^ Library Validation and Cluster Density Optimization.

13. Kebschull JM, Garcia da Silva P, Reid AP, Peikon ID, Albeanu DF, Zador AM. 2016. High-Throughput Mapping of Single-Neuron Projections by Sequencing of Barcoded RNA. Neuron 91: 975–987.

14. Kinde I, Wu J, Papadopoulos N, Kinzler KW, Vogelstein B. 2011. Detection and quantification of rare mutations with massively parallel sequencing. Proc Natl Acad Sci U S A 108: 9530–5.

15. Kivioja T, Vaharautio A, Karlsson K, Bonke M, Enge M, Linnarsson S, Taipale J. 2011. Counting absolute numbers of molecules using unique molecular identifiers. Nat Methods 9: 72.

16. Koike-Yusa H, Li Y, Tan E-P, Velasco-Herrera MDC, Yusa K. 2014. Genome-wide recessive genetic screening in mammalian cells with a lentiviral CRISPR-guide RNA library. Nat Biotechnol 32: 267–273.

17. Langmead B, Salzberg SL. 2012. Fast gapped-read alignment with Bowtie 2. Nat Methods 9: 357–359.

18. Martin M. 2011. Cutadapt removes adapter sequences from high-throughput sequencing reads. EMBnet.journal 17: 10–12.

19. McKenna A, Findlay GM, Gagnon JA, Horwitz MS, Schier AF, Shendure J. 2016. Whole-organism lineage tracing by combinatorial and cumulative genome editing. Science (80- ) 353: aaf7907.

20. Peikon ID, Kebschull JM, Vagin V V, Ravens DI, Brouzes E, Correa IR, Bressan D, Zador A. 2017. Using high-throughput barcode sequencing to efficiently map connectomes. bioRxiv 099093.

21. Ponts N, Harris EY, Prudhomme J, Wick I, Eckhardt-Ludka C, Hicks GR, Hardiman G, Lonardi S, Le Roch KG. 2010. Nucleosome landscape and control of transcription in the human malaria parasite. Genome Res 20: 228–38.

22. Rodriguez-Barrueco R, Marshall N, Silva JM. 2013. Pooled shRNA screenings: experimental approach. Methods Mol Biol 980: 353–70.

23. Shalem O, Sanjana NE, Hartenian E, Shi X, Scott DA, Mikkelsen TS, Heckl D, Ebert BL, Root DE, Doench JG, et al. 2014. Genome-Scale CRISPR-Cas9 Knockout Screening in Human Cells. Science (80- ) 343: 84–87.

24. Shalem O, Sanjana NE, Zhang F. 2015. High-throughput functional genomics using CRISPR– Cas9. Nat Rev Genet 16: 299–311.

25. Sims D, Mendes-Pereira AM, Frankum J, Burgess D, Cerone M-A, Lombardelli C, Mitsopoulos C, Hakas J, Murugaesu N, Isacke CM, et al. 2011. High-throughput RNA interference screening using pooled shRNA libraries and next generation sequencing. Genome Biol 12: R104.

26. Smith AM, Heisler LE, Mellor J, Kaper F, Thompson MJ, Chee M, Roth FP, Giaever G, Nislow C. 2009. Quantitative phenotyping via deep barcode sequencing. Genome Res 19: 1836–42.

27. Strezoska Z, Licon A, Haimes J, Spayd KJ, Patel KM, Sullivan K, Jastrzebski K, Simpson KJ, Leake D, van Brabant Smith A, et al. 2012. Optimized PCR conditions and increased shRNA fold representation improve reproducibility of pooled shRNA screens. ed. K.T. Jeang. PLoS One 7: e42341.

28. Taylor DL, Walters WA, Lennon NJ, Bochicchio J, Krohn A, Caporaso JG, Pennanen T. 2016. Accurate Estimation of Fungal Diversity and Abundance through Improved Lineage-Specific Primers Optimized for Illumina Amplicon Sequencing. Appl Environ Microbiol 82: 7217–7226.

29. van Opijnen T, Camilli A. 2013. Transposon insertion sequencing: a new tool for systems-level analysis of microorganisms. Nat Rev Microbiol 11: 435–442.

30. Wang T, Wei JJ, Sabatini DM, Lander ES. 2014. Genetic Screens in Human Cells Using the CRISPR-Cas9 System. Science (80- ) 343: 80–84.

31. Woese CR, Magrum LJ, Gupta R, Siegel RB, Stahl DA, Kop J, Crawford N, Brosius R, Gutell R, Hogan JJ, et al. 1980. Secondary structure model for bacterial 16S ribosomal RNA: phylogenetic, enzymatic and chemical evidence. Nucleic Acids Res 8: 2275–2294.

32. Woese CR, 2012a.) A framework for human microbiome research. Nature 486: 215–21.

33. Woese CR, 2012b.) Evaluation of 16S rDNA-based community profiling for human microbiome research. PLoS One 7: e39315.

